# The conserved β-hairpin of the SUI1 domain is a dual-function structural module governing translation initiation and ribosome recycling in yeast

**DOI:** 10.64898/2026.08.14.744980

**Authors:** Kseniya A. Zamyatnina, Valery N. Urakov, Inna A. Volynkina, Elena A. Stolboushkina, Evgeny S. Gerasimov, Leonid M. Kats, Vitaly V. Kushnirov, Piotr A. Kamenski, Sergey E. Dmitriev

## Abstract

Most eukaryotic mRNAs encode a single functional polypeptide. Following translation termination, both the large and small ribosomal subunits are typically released from the mRNA by ribosome recycling factors. However, after translating short upstream open reading frames (uORFs) within the 5’ untranslated regions (UTRs), ribosomes can remain associated with the mRNA and reinitiate translation. This process is regulated by the heterodimer MCTS1•DENR (Tma20p•Tma22p in yeast). DENR/Tma22p harbors a SUI1 domain, structurally homologous to the translation initiation factor eIF1/Sui1p, which features a conserved, positively charged β-hairpin loop critical for eIF1 function. Despite this structural similarity, the functional significance of specific elements within DENR/Tma22p remains unexplored. Here, we used *in vivo* reporter assays in *Saccharomyces cerevisiae* to quantify reinitiation efficiency following translation of either a short uORF (in the 5’ UTR) or a full-length coding sequence (in the 3’ UTR). Systematic analysis of single, double, and triple deletions of *TMA20*, *TMA22*, and *TMA64* (a homolog of Tma20p•Tma22p) revealed that the Tma20p•Tma22p complex exerts a dominant role over Tma64p in modulating reinitiation, while exhibiting functional interplay between the two factors. Using knockout strains complemented with Tma22p variants, we further demonstrated that the positively charged residues of the β-hairpin loop 1 are essential for Tma22p recycling activity. Unexpectedly, deletion of the entire SUI1 domain was less deleterious, and eIF1/Sui1p was able to partially substitute for the SUI1 domain of Tma22p within a chimeric protein context. Our findings establish the β-hairpin loop 1 of the DENR/Tma22p SUI1 domain as a critical determinant for ribosome recycling and reinitiation, and raise the question of whether MCTS1/Tma20p can promiscuously operate with either DENR/Tma22p or eIF1/Sui1p – two specialized factors that evolved from a common structural scaffold to govern distinct steps in the translation cycle.

## Introduction

In eukaryotic cells, the majority of mRNAs are monocistronic, encoding a single polypeptide. Following translation termination, the ribosomal complex must be disassembled into its constituent subunits to be reused in subsequent rounds of translation (Hellen, 2018; Tahmasebinia & Wu, 2025; Young & Guydosh, 2022). This essential process, known as ribosome recycling, occurs in two main stages. The first step, the dissociation of the 60S subunit from the post-termination complex, is catalyzed by the ATP-binding cassette protein ABCE1 (Rli1p in yeast) (Pisarev et al, 2010; Shoemaker & Green, 2011; Young et al, 2015), which is also involved in translation initiation and termination. The subsequent step, which includes the dissociation of the 40S subunit from the mRNA and deacylated P-site bound tRNA, is facilitated by specialized recycling factors. In mammals, this function is majorly performed by the heterodimer MCTS1•DENR, while in yeast *Saccharomyces cerevisiae*, this role is fulfilled by the respective orthologs, Tma20p•Tma22p (Skabkin et al, 2010; Young et al, 2018; Young et al, 2021). Alternative pathways of 40S recycling have been reported, involving either initiation factors eIF1, eIF1A, eIF3, and eIF3j (Pisarev et al, 2007; Pisarev et al, 2010), or eIF2D/Tma64p (Dmitriev et al, 2010; Skabkin et al, 2010; Young et al, 2018). The latter, although clearly implemented in or contributing to 40S recycling in many models and experimental setups (Bohlen et al, 2023; Jendruchova et al, 2024; Skabkin et al, 2010; Skabkin et al, 2013; Vasudevan et al, 2020; Young et al, 2018) and exhibiting a negative genetic interaction with Tma20p (Costanzo et al, 2016), has recently been challenged as a key factor operating at stop codons (Gaikwad et al, 2021; Jendruchova et al, 2024; Meurs et al, 2025; Young et al, 2021), and has finally been implicated in 40S ribosomal subunit recycling during intrinsic ribosome destabilization (Ichihara et al, 2025). The proteins MCTS1, DENR, and eIF2D are widely associated with many human diseases, such as cancer, immune system failure, and neurological disorders (Bohlen et al, 2023; Shyrokova et al, 2021; Tahmasebinia & Wu, 2025; Zamyatnina, 2025).

A critical physiological context where ribosome recycling modulates mRNA translation efficiency is reinitiation. This process occurs when a ribosome that has terminated translation at a stop codon does not dissociate from the mRNA but instead resumes scanning and initiates translation at a downstream start codon (Dever et al, 2023; Gunisova et al, 2018; Jackson et al, 2010; Young & Guydosh, 2022). Reinitiation is a common mechanism for regulating protein synthesis, often occurring after the translation of short upstream open reading frames (uORFs) found in the 5’ untranslated regions (UTRs) of many eukaryotic, especially mammalian, mRNAs (Dever et al, 2023; Gunisova et al, 2018; Hellen, 2018). In this context, an inefficient or delayed ribosome recycling at the stop codon of an uORF usually upregulates translation of the main coding region (Jendruchova et al, 2024; Makeeva et al, 2019; Skabkin et al, 2013; Young et al, 2015; Young et al, 2018). Reinitiation can also occur on bicistronic mRNAs (having two long translons encoding functional proteins), both viral (Sorokin et al, 2021; Zinoviev et al, 2015) and rarely occurring cellular ones (Andreev & Shatsky, 2025; Dmitriev et al, 2007). At regular monocistronic mRNAs, the post-termination complex is preferentially recycled to enable ribosomes to participate in a new round of translation, including reinitiation on the same mRNA via a process termed CLAR (closed-loop assisted reinitiation, see (Alekhina et al, 2020) and references therein). Here, defective recycling leads to aberrant translation of the 3’ UTR (Young et al, 2015; Young et al, 2018) and can generate toxic polypeptide products (Tahmasebinia & Wu, 2025).

Interestingly, the structural organization of the post-termination complex resembles that of the translation initiation complex: in both cases, the 40S subunit is bound to an mRNA with a vacant A site and a tRNA positioned in the P site. The key difference lies in the identity of the tRNA – a deacylated tRNA in recycling versus the initiator Met-tRNA_i_^Met^ in initiation complexes. This similarity, supported by structural data (see below), is underscored by the fact that in reconstituted translation systems, MCTS1•DENR and eIF2D were shown to both stabilize cognate tRNAs in the P site during initiation on certain viral and leaderless mRNAs and destabilize deacylated tRNAs during recycling (Dmitriev et al, 2010; Skabkin et al, 2010). This duality has made it challenging to define their precise physiological functions, especially in the context of reinitiation.

In particular, studies in drosophila and mammalian systems have revealed (Bohlen et al, 2020; Castelo-Szekely et al, 2019; Chen et al, 2022; Haas et al, 2016; Meurs et al, 2025; Schleich et al, 2017; Schleich et al, 2014; Vasudevan et al, 2020) that knockouts (KOs) and knockdowns of *MCTS1*, *DENR*, and *EIF2D* produce effects contrary to those observed in yeast. These effects have been validated and repeatedly confirmed on both reporter and natural mRNAs, and were interpreted to suggest that the MCTS1•DENR complex (and likely eIF2D) can facilitate the delivery of Met-tRNA_i_ ^Met^ during reinitiation, thereby serving as a surrogate for the canonical initiation factor eIF2. This hypothesis aligns with early data obtained in reconstituted mammalian systems (Dmitriev et al, 2010; Skabkin et al, 2010; Skabkin et al, 2013), but largely contradicts results in yeast, where deletion of the corresponding genes led to increased reinitiation, both after full-length coding regions and following short uORFs (Gaikwad et al, 2021; Jendruchova et al, 2024; Makeeva et al, 2019; Young et al, 2018; Young et al, 2021). This discrepancy can be explained by a principal difference in ribosome recycling mechanisms employed by yeast and animals, different reporter systems used in these studies, or both. Thus, despite their similar functional context, a unified model of the molecular mechanisms governing their activity remains elusive.

The dimer MCTS1•DENR/Tma20p•Tma22p and eIF2D/Tma64p share a common domain architecture (Ahmed et al, 2018; Lomakin et al, 2020; Lomakin et al, 2019; Lomakin et al, 2017; Vaidya et al, 2017; Weisser et al, 2017) (Fig. 1a). Both protein families contain a DUF1947-PUA superdomain (MCTS1/Tma20p or the N-terminal part of eIF2D/Tma64p, respectively) involved in 40S binding and stabilization of the P-site bound tRNA (Lomakin et al, 2017; Weisser et al, 2017). In addition, eIF2D possesses Winged Helix and SWIB/MDM2 domains (Vaidya et al, 2017; Weisser et al, 2017), while DENR instead has a zinc-binding minidomain (ZnBD) that interacts with MCTS1 (Ahmed et al, 2018; Lomakin et al, 2019). A defining feature is the presence of a C-terminal SUI1 domain, which is homologous to the SUI1 domain of the translation initiation factor eIF1/Sui1p (Fletcher et al, 1999; Reibarkh et al, 2008).

**Figure 1.**
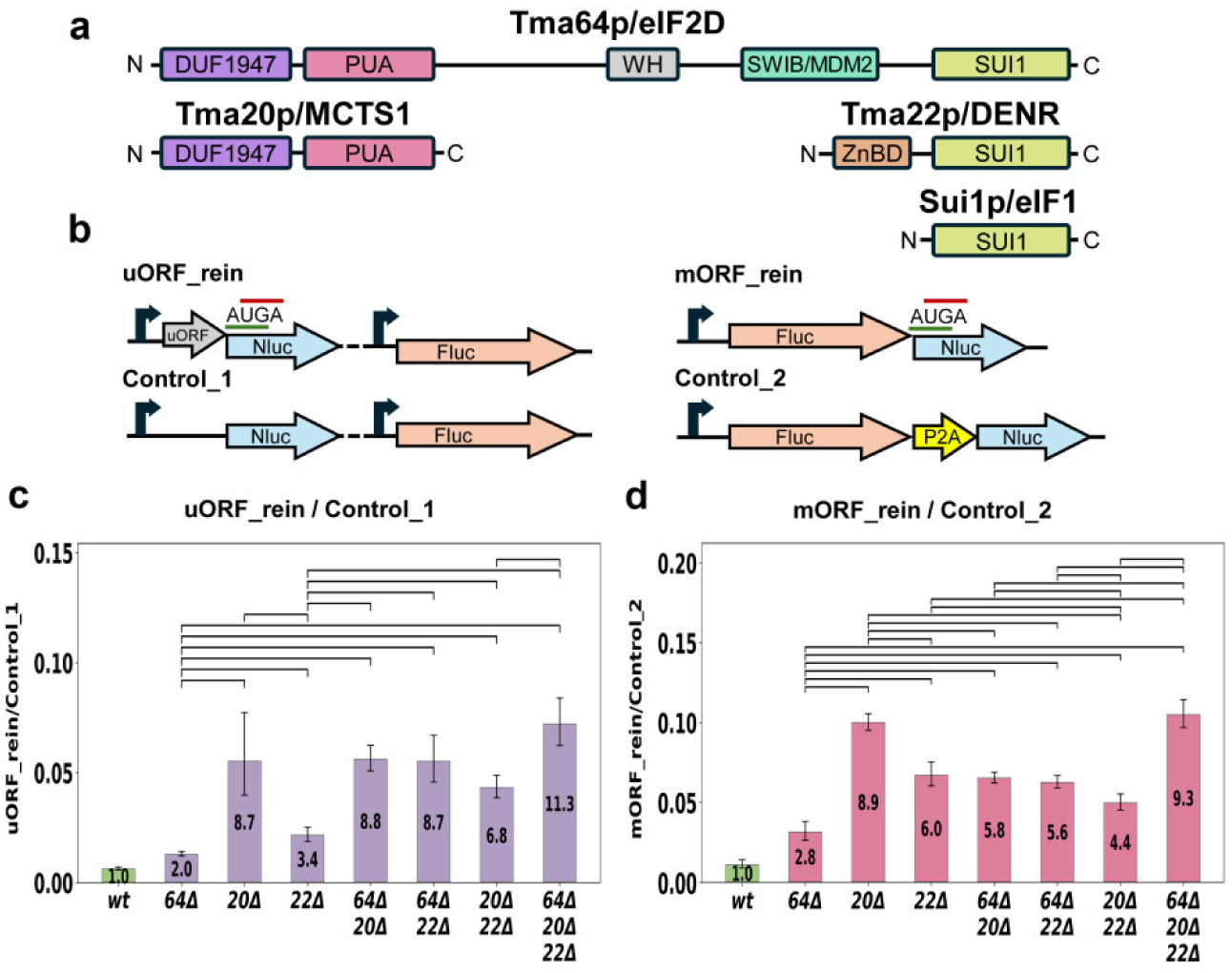
Yeast strains lacking 40S recycling factors exhibit elevated translation reinitiation rates. *in vivo*, **as revealed by reporter expression analysis.** (**a**) Domain organization of yeast and human Tma64p/eIF2D, Tma20p/MCTS1, and Tma22p/DENR. (**b**) Schematic representation of the reporter constructs used in this study. Nluc activity from the uORF_rein and mORF_rein constructs reflects reinitiation activity, whereas Fluc activity serves as a normalization control. Control_1 and Control_2 constructs were included for additional normalization, as the Nluc/Fluc ratio can vary between strains. Promoters are indicated by arrows; AUGA represents a stop-start reinitiation sequence; P2A denotes a StopGo (2A-peptide) sequence. (**c**, **d**) Relative translation reinitiation rates in *wt* and single-, double-, or triple-knockout strains (*tma20Δ*, *tma22Δ*, and *tma64Δ*). Rates were calculated as the normalized Nluc activity (Nluc/Fluc ratio) from the uORF_rein (c) and mORF_rein (d) constructs, divided by the corresponding Nluc/Fluc ratio from the “Control” constructs. Numbers inside the bars indicate the fold increase in reinitiation rates relative to the *wt* strain. Brackets denote pairs of mutants with statistically significant differences (*p* < 0.001). All mutants differ significantly from the *wt* strain (not indicated in the graph). Experiments were performed in at least three biological replicates. Mean values and corresponding 95% confidence intervals are shown.

The eIF1 SUI1 domain plays a pivotal role in start codon selection (for review, see (Hinnebusch, 2014)). Structural studies of eukaryotic translation pre-initiation complexes have revealed that the eIF1 SUI1 domain binds adjacent to the P site, with its conserved positively charged β-hairpin loop 1 protruding toward the mRNA-tRNA_i_ codon-anticodon base pairs (Hussain et al, 2014; Llacer et al, 2015; Lomakin et al, 2003; Lomakin & Steitz, 2013; Rabl et al, 2011; Weisser et al, 2013). Biochemical and genetic data indicate that basic residues within this loop are involved in stable 40S binding and regulate the transition of Met-tRNA_i_ from the P_OUT_ to the P_IN_ state upon AUG recognition (Martin-Marcos et al, 2013). The current model suggests that β-hairpin 1 alternates between two states, tracking the Met-tRNA_i_ anticodon stem-loop during the transition of the 48S complex from an open to a closed conformation, and substantially contributes to the destabilization and subsequent dissociation of eIF1 from the 40S subunit upon AUG codon recognition (reviewed in (Hinnebusch, 2017)).

Intriguingly, structural studies of human DENR or eIF2D bound to the 40S ribosomal subunit have shown that the SUI1 domains of both proteins occupy the same binding site as eIF1, with the basic β-hairpin loop 1 directed toward mRNA cleft of the P site (Lomakin et al, 2017; Weisser et al, 2017). One may assume that this conserved structural element has been functionally repurposed for ribosome recycling and/or translation reinitiation. However, no genetic or biochemical data on its importance for these processes have been obtained to support this idea.

In this study, we employed *in vivo* reporter assays in *S. cerevisiae* to dissect the functional requirements of the Tma22p SUI1 domain and to delineate the relative contributions of Tma64p and the Tma20p•Tma22p heterodimer in regulating reinitiation. Using KO strains complemented with specific Tma22p derivatives, including those with amino acid substitutions in the β-hairpin loop 1, domain deletions, and SUI1 domain replacement with that of Sui1p, and systematically analyzing single, double, and triple KO strains, we demonstrate that the Tma22p SUI1 domain with positively charged β-hairpin loop 1 is essential for its recycling activity. These findings provide new mechanistic insights into how a conserved structural scaffold has been adapted to govern distinct steps in the translation cycle.

## Results

### A Reporter System for Monitoring Translation Reinitiation Efficiency

We previously developed a set of luciferase-based reporters for the quantitative analysis of translation reinitiation rates in a yeast cell-free system (Young et al, 2018). We found that an mRNA containing a relatively short, simple CA-rich leader with a single 5-codon uORF, terminated just upstream of the firefly luciferase (*Fluc*) coding region (CDS), shows a pronounced increase in Fluc activity (indicating reinitiation) in ribosome recycling factor-deficient strains. Specifically, Fluc activity was 5-6-fold higher in *tma20Δtma64Δ* and *tma22Δtma64Δ* double KO strains compared to the wild-type (*wt*) strain. We also established that in a bicistronic construct, where the *Fluc* CDS followed a full-length protein-encoding ORF, the reporter exhibited even higher (7-8-fold) activation in the mutant strains (Young et al, 2018). These two types of reporters can be regarded as models of “legitimate”, regulatory reinitiation (in the 5′ UTR after reading an uORF) or “illegitimate”, undesirable reinitiation (in the 3′ UTR after translating the CDS), respectively. However, such reporters have never been used *in vivo*, nor have their activities been analyzed beyond the double KO strains.

Thus, we constructed a set of bicistronic reporter plasmids based on the centromeric pCM189 vector (Fig. 1b, Suppl. Data 1). The “uORF_rein” construct produces two independent mRNAs: the first contains the Nano-luciferase (*Nluc*) CDS preceded by a 5-codon uORF (with no spacer between them), and the second encodes Fluc as a normalization reporter. This was complemented by “Control_1”, which produces similar transcripts, except that the *Nluc* transcript lacked the uORF. In contrast, the “mORF_rein” construct produces a single bicistronic mRNA where the *Nluc* CDS immediately followed the full-length *Fluc* CDS (thus representing an ORF located within the 3′ UTR). This construct was complemented by “Control_2”, where Fluc and Nluc were produced from the same ORF, separated by a sequence encoding a 2A-peptide (StopGo element), to produce equimolar amounts of the two reporters. It should be noted that all encoded mRNAs had relatively short and simple 5′ UTRs, similar to those used in the previous study (Young et al, 2018).

Therefore, in both cases, Nluc activity reflected the rate of reinitiation that occurred after translation of either a short uORF in the 5′ UTR or a long “main ORF” of an mRNA. In all experiments described below, relative reinitiation rates were evaluated as the Nluc/Fluc activity ratio for a “rein” construct, normalized to the corresponding ratio from its respective “Control” construct. To assess the effects of mutation(s) in the studied strains, these values were further normalized to those obtained in the *wt* strain. This design excluded possible differential effects of the studied mutations on reporter transcription, luciferase activities, or other assay components. Furthermore, it should render the results independent of most other non-specific translation reprogramming and perturbations, including those observed previously in *TMA* KO strains (Gaikwad et al, 2021).

### The Tma20p•Tma22p complex plays a dominant role over Tma64p in modulating reinitiation

First, to dissect the relative contributions of Tma64p and the Tma20p•Tma22p heterodimer to the suppression of reinitiation activity, we assayed our reporters in strains harboring single (*tma64Δ*, *tma20Δ*, *tma22Δ*), double (*tma64Δtma20Δ*, *tma64Δtma22Δ*, *tma20Δtma22Δ*), and triple (*tma64Δtma20Δtma22Δ*) KOs.

In accordance with results previously obtained in cell-free systems (Young et al, 2018), the *tma64Δtma20Δ* and *tma64Δtma22Δ* double KO strains exhibited a substantially elevated reinitiation rate for both reporter systems (Fig. 1c,d). This most likely reflects a ribosome recycling deficiency due to the absence of both the Tma20p•Tma22p and Tma64p 40S recycling factors. The effects were roughly equivalent for the two mutants. The third double KO strain, *tma20Δtma22Δ*, which lacks Tma20p•Tma22p but possesses Tma64p, showed a similar, albeit somewhat lower, level of reinitiation. This suggests a minor but present contribution of Tma64p to the process. This conclusion was further supported by the triple KO strain, *tma64Δtma20Δtma22Δ*, which exhibited the highest level of reinitiation for both reporter systems.

Notably, single gene KOs resulted in differential effects on reinitiation rates. Deletion of *TMA20* produced the most prominent elevation of reinitiation in both reporter systems, while the *TMA64* KO had the lowest, albeit statistically significant, effect (consistent with previous reports (Gaikwad et al, 2021; Jendruchova et al, 2024; Meurs et al, 2025; Young et al, 2021) and the recently identified role of eIF2D in a distinct process (Ichihara et al, 2025)). Intriguingly, in our hands, the single *TMA22* KO produced a considerably lesser stimulation of reinitiation than the deletion of *TMA20*, which encodes its partner in the Tma20p•Tma22p heterodimer. This unexpected result was reproduced in both the uORF_rein and mORF_rein experimental systems (although it was more prominent in the former one), suggesting possibilities of Tma22p replaceability and promiscuous Tma20p activity, which we discuss below.

We also noted that the extent to which reinitiation rates were elevated slightly differed between our two reporter systems (compare Fig. 1c and Fig. 1d). Specifically, single gene deletions produced somewhat higher effects with the mORF_rein construct, whereas double and triple KOs resulted in greater stimulation of reinitiation activity with the uORF_rein construct. Although difficult to interpret due to fundamental differences in reporter design (see also Discussion), this could reflect a more versatile mechanism of ribosome recycling or reinitiation after translation of short uORF compared to that following the translation of a long main ORF.

In summary, these results lead us to conclude that the Tma20p•Tma22p heterodimer, and especially its Tma20p component, make a greater contribution to the suppression of reinitiation activity after both uORF and main ORF (CDS) translation.

### Disruption of functional elements in Tma22p elevates translation reinitiation rate in mutant yeast strains

Based on the above results, we selected the *tma64Δtma22Δ* strain as the foundation for further study of Tma22p. To identify critical functional elements within Tma22p, we generated a number of strains expressing the *TMA22* gene, either in its *wt* form or with specific mutations (Fig. 2a). In particular, we created two strains producing Tma22p variants with point substitutions in the positively charged β-hairpin loop 1 of the SUI1 domain (KRK108AEA and K110E) and two mutants with large Tma22p deletions: lacking the ZnBD (L2-E90del) or SUI1 (L95*) domains (Fig. 2b). These strains were transformed with the reporter plasmids, and reinitiation efficiencies were measured as described above (Fig. 2c,d).

**Figure 2.**
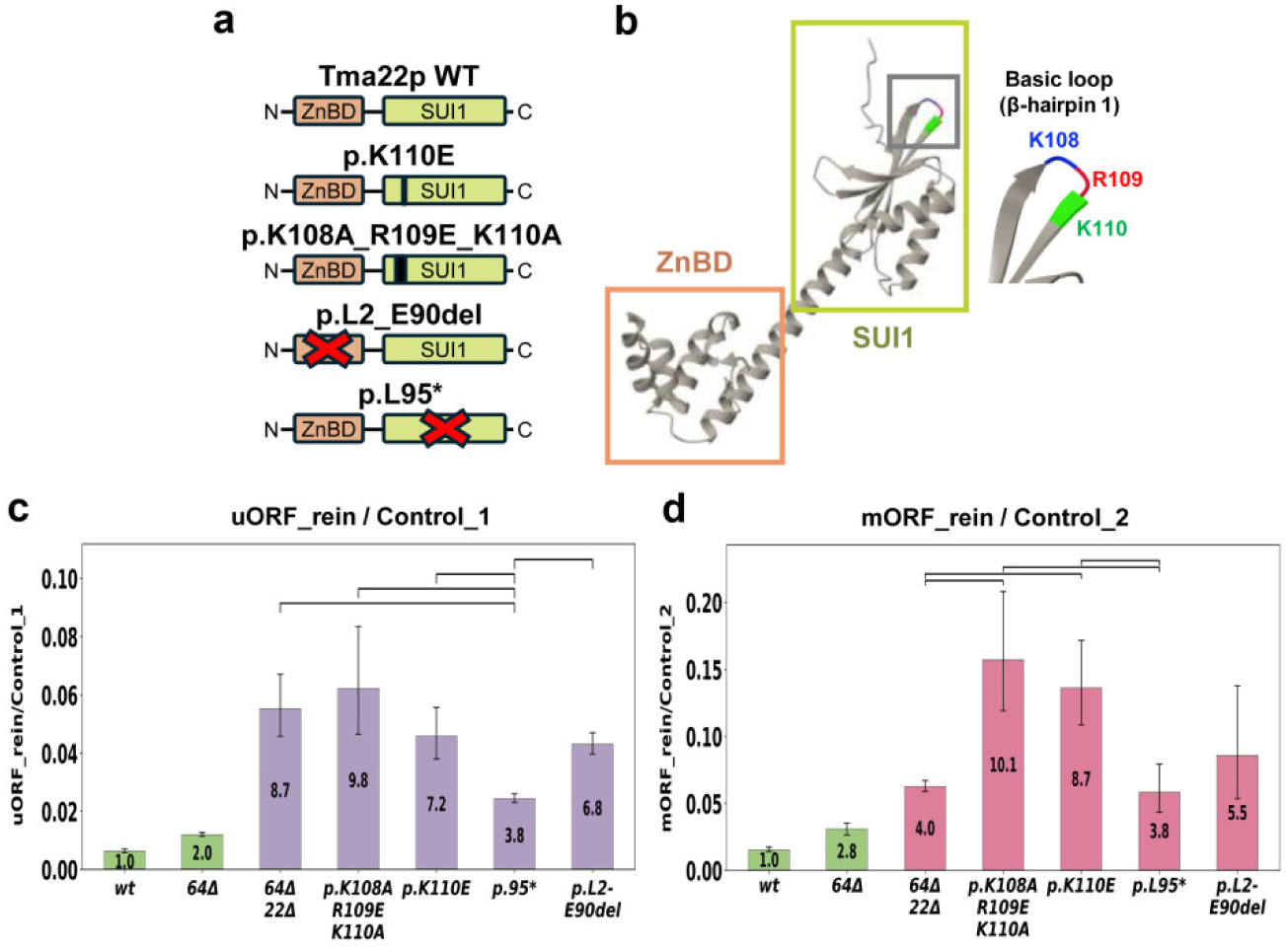
Disruption of functional elements in Tma22p elevates translation reinitiation rate in mutant yeast strains. (**a**) Domain organization of Tma22p and its variants. Amino acid substitutions are marked by black lines, and complete domain deletions are indicated by red crosses. (**b**) AlphaFold-predicted 3D structure of Tma22p with structured domains highlighted and the basic β-hairpin loop 1 shown in a close-up view. (**c**, **d**) Relative translation reinitiation rates in *wt*, *tma64Δ*, *tma64Δtma22Δ*, and *tma64Δ*-based strains with *TMA22* variants introduced in place of the *wt* gene. Indications are the same as in Fig. 1. Brackets denote statistically significant differences in reinitiation rates (*p* < 0.001). All mutants showed significant differences from the *wt* and *tma64Δ* strains (not indicated). Experiments were performed in at least three biological replicates. Mean values and corresponding 95% confidence intervals are shown.

Neither of the Tma22p derivatives was able to completely rescue the *tma64Δtma22Δ* phenotype (i.e., to reduce the reinitiation rate of either reporter to that observed in the *wt* or *tma64Δ* strains expressing the native *TMA22* gene). However, the different variants yielded distinct reinitiation levels. Both point substitutions (KRK108AEA and K110E) in the positively charged β-hairpin loop 1 of the SUI1 domain resulted in the complete loss of Tma22p recycling activity. Specifically, reinitiation rates observed in strains expressing KRK108AEA or K110E versions of Tma22p were roughly similar or even slightly higher than those observed in the strain with *TMA22* deletion. We note that the effect of the triple substitution (KRK108AEA) was likely more pronounced, especially in the context of the mORF_rein construct (Fig. 2d). Although these effects were not statistically significant, this suggests that the binding of a non-functional factor to the ribosome may be more deleterious than the complete absence of the factor.

Even more intriguing results were obtained using strains producing Tma22p with domain deletions. While removing the N-terminal (ZnBD, Tma20p-interacting) domain resulted in effects similar to those observed in the parental *tma64Δtma22Δ* strain (suggesting inability of Tma22p to function without Tma20p binding), the strain lacking the C-terminal (SUI1) domain produced significantly lower reinitiation rates in both reporter systems. The latter result was unexpected, especially in light of the above dramatic effects of point substitutions in this domain. This could be explained by the involvement of other participants (e.g., Sui1p/eIF1) that are able to substitute the Tma22p SUI1-domain activity, as discussed below. It should also be noted that the Tma20p-interacting ZnBD likely protects Tma20p from destabilization, a phenomenon known for animal MCTS1 in the absence of DENR (see (Ahmed et al, 2018) and references therein). This may explain why Tma20p alone did not exhibit similar activity in the *tma64Δtma22Δ* strain.

Together, these results highlight the critical importance of the positively charged β-hairpin loop 1 of the SUI1 domain and the ZnBD domain of Tma22p, but also suggest the presence of alternative SUI1-domain-like activity involved in ribosome recycling in yeast cells.

### The Tma22p-Sui1p chimeric protein partially restores ribosome recycling in the *tma22Δ* strain

To further investigate whether the SUI1 domain of Sui1p/eIF1 can substitute its counterpart in Tma22p protein function during 40S recycling, we constructed an integrative plasmid expressing a chimeric protein where the N-terminal ZnBD domain of Tma22p (amino acid residues 1-77) was fused to the SUI1 domain of Sui1p (amino acid residues 27-108) (Fig. 3a). This construct was integrated into W303 *wt* and *tma22Δ* strains. The resulting four strains (Fig. 3b) were assayed with the same two reporters. It should be noted that, due to technical reasons, we used the W303 background instead of BY4741 in these experiments, and all strains maintained an intact *TMA64* gene.

**Figure 3.**
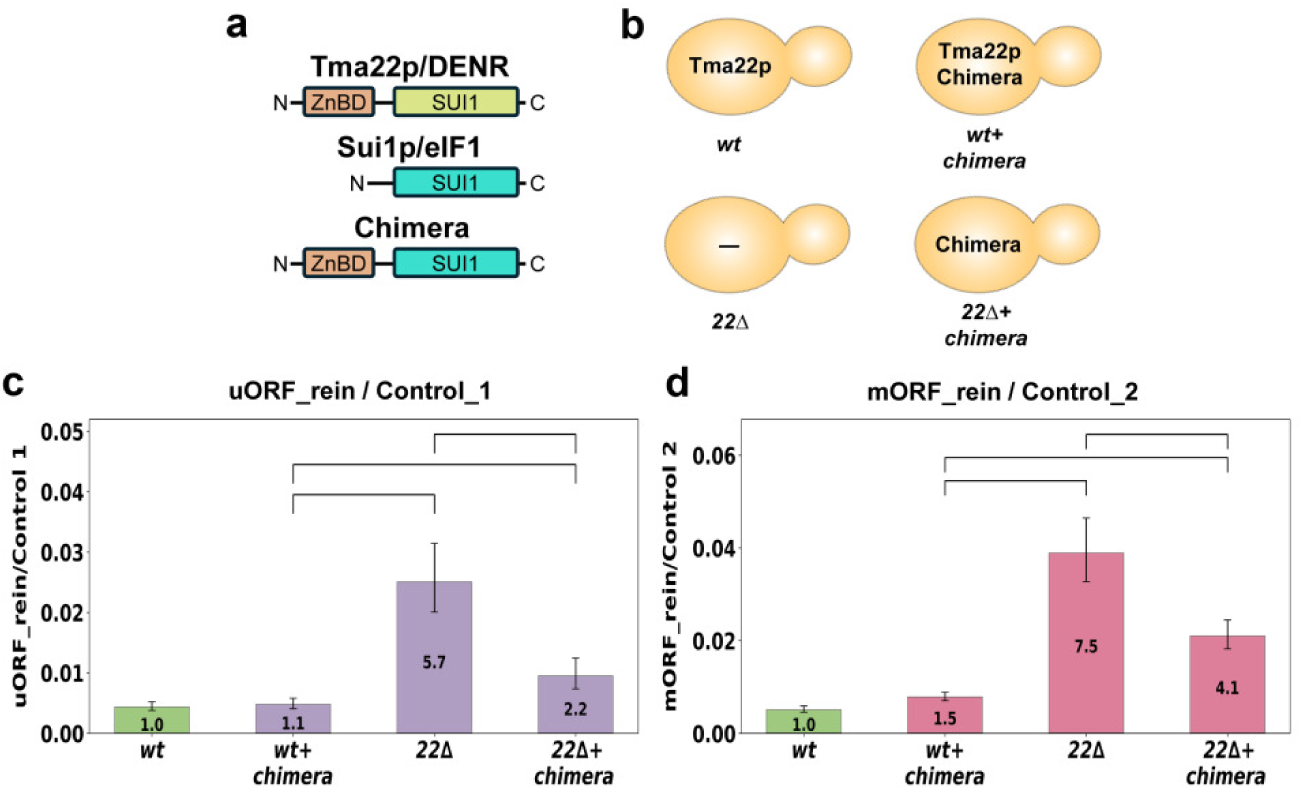
The Tma22p-Sui1p chimeric protein partially restores reinitiation suppression in the *tma22Δ* strain. (**a**) Domain organization of Tma22p, Sui1p, and the Tma22p-Sui1p chimera. (**b**) The W303-based strains used in the analyses. The chimera was expressed from an integrative plasmid in either *wt* or a *tma22Δ* background, producing the indicated proteins. (**c**, **d**) Relative translation reinitiation rates in the analyzed strains. Symbols and color-coding are the same as in Fig. 1. Brackets denote statistically significant differences in reinitiation rates (*p* < 0.001). Experiments were performed in at least three biological replicates. Mean values and corresponding 95% confidence intervals are shown.

In the *wt* strain expressing the chimeric protein (*wt+chimera*), reinitiation after mORF translation showed a minor, statistically insignificant increase compared to the *wt* strain expressing endogenous Tma22p (Fig. 3c,d). This suggests that the Chimera either inefficiently competes with endogenous Tma22p for ribosome binding or is as active as the *wt* protein. As expected, deletion of *TMA22* resulted in elevated reinitiation levels in both reporters. However, the presence of the Chimera protein partially rescued this phenotype of the *tma22Δ* strain. This effect was statistically significant for both uORF_rein and mORF_rein reporter systems.

These data suggest that the SUI1 domain of yeast eIF1 can partially substitute the corresponding part of the Tma22p/DENR protein in 40S recycling.

## Discussion

In this study, we have used a combination of reporter assays and targeted mutagenesis in *S. cerevisiae* to delineate the roles of the Tma20p•Tma22p heterodimer and Tma64p in regulating ribosome recycling and reinitiation, and to define the functional requirements of the Tma22p SUI1 domain. As mutations in the *TMA* genes have been previously shown to reprogram translation in cells to some extent (Gaikwad et al, 2021), we used a sophisticated two-step normalization approach to minimize potential non-specific effects of mutations on reporter activity. We also used two reporter systems to separately study reinitiation in the 5′ and 3′ UTRs (uORF_rein and mORF_rein, respectively).

First, we found that Tma20p•Tma22p makes a major contribution to the suppression of reinitiation activity over Tma64p after both uORF and main ORF translation, i.e., in both 5′ and 3′ UTRs. Our results from single gene deletions also suggest that the Tma20p component of the heterodimer is likely more important for this activity than its partner Tma22p, suggesting that some other components of the translational machinery (e.g., eIF1/Sui1p) can partially compensate for the activity of the latter. Second, we showed that positively charged amino acid residues in the β-hairpin loop 1 of the SUI1 domain of Tma22p are critically important for its function and that Tma22p interaction with Tma20p *via* the ZnBD domain is indispensable. In contrast, deletion of the complete SUI1 domain had a much less pronounced effect on proper ribosome recycling and reinitiation in cells, suggesting its compensation by other participants, such as eIF1/Sui1p. Third, we demonstrated that the SUI1 domain from Sui1p/eIF1 is likely able to support 40S recycling activity when it replaces the original SUI1 domain in Tma22p. Together, these results suggest that, at least under some conditions, MCTS1/Tma20p can promiscuously operate with eIF1/Sui1p instead of DENR/Tma22p.

Our conclusion that the Tma20p•Tma22p complex exerts a dominant role over Tma64p in modulating reinitiation fully aligns with a number of recent reports (Gaikwad et al, 2021; Jendruchova et al, 2024; Makeeva et al, 2019; Meurs et al, 2025; Young et al, 2021). In particular, using systems biology approaches such as ribosome profiling or TCP-seq, studies from the Hinnebusch and Guydosh laboratories have shown (Gaikwad et al, 2021; Young et al, 2021) that the *tma64Δ* single mutant displays minimal or no signs of impaired 40S recycling. However, focused analysis of specific mRNA cohorts revealed that their translation exhibited qualitatively comparable, albeit much less pronounced (an order of magnitude lower), changes in the *tma64Δ* strain compared to the *tma64Δtma20Δ* strain (Gaikwad et al, 2021). Next, in a recent study by the Valášek and Hinnebusch groups (Jendruchova et al, 2024), single, double, and triple KO strains (similar to those used in our study) were systematically assayed using a *LacZ* reporter containing a *GCN4*-derived 5′ UTR harboring a single short uORF. Although mostly focused on the effects of the uORF penultimate codon, this study clearly showed that Tma64p poorly substitutes for the Tma20p•Tma22p heterodimer in suppressing reinitiation. However, in many cases, knocking out *TMA64* in addition to *TMA20*, *TMA22*, or both caused a stronger phenotype. All these results correlate well with our earlier analysis of ribosome profiling data (Makeeva et al, 2019), focusing on *GCN4* and *MATa2* mRNAs, and with our current observations.

Nevertheless, we would like to note that the relative contribution of deleting a particular gene may be determined not only by the protein’s function but also by its expression level (protein abundance) under specific experimental conditions. For example, according to the SGD database (https://yeastgenome.org/), Tma64p abundance is approximately half that of Tma20p•Tma22p (∼5,000 molecules/cell on average for Tma64p vs. ∼10,000 for both Tma20p and Tma22p). This protein may have a specific function, as *TMA64* gene expression is highly upregulated during a specific stage of sporulation (see expression profiles in (Brar et al, 2012; Primig et al, 2000) or other relevant studies) and in some other contexts. It would be of interest to analyze the differential contribution of Tma proteins under these conditions. When comparing the effects of individual KOs, the potential influence of altered expression of genes adjacent to the knocked-out genes must also be considered (Egorov et al, 2021; Makeeva et al, 2019). However, taken together with recent findings from mammalian systems (Meurs et al, 2025), these results allow us to conclude that the Tma20p•Tma22p heterodimer (MCTS1/DENR in mammals) is likely a major source of 40S recycling activity at stop codons of both uORFs and main ORFs. In contrast, Tma64p/eIF2D likely has a distinct primary function but can participate in this process under conditions of Tma22p/DENR and, especially, Tma20p/MCTS1 depletion (which is also reflected in a negative genetic interaction between the *TMA64* and *TMA20* genes found in genome-wide screenings, such as (Costanzo et al, 2016)). Consistent with this, eIF2D has been recently implicated in 40S recycling within the CDS during intrinsic ribosome destabilization (Ichihara et al, 2025).

Intriguingly, we also found that single *TMA22* deletion causes a significantly weaker effect on reinitiation than that of *TMA20*, especially in the context of uORFs, suggesting a compensatory activity in yeast cells. This was unexpected, as previous studies using such strains did not report a substantial difference between the phenotypes of single *tma20Δ* and *tma22Δ* mutants. However, re-examination of the data from previous studies (Gaikwad et al, 2021; Young et al, 2018; Young et al, 2021) with this new focus revealed that the increases in ribosomal peaks at or near stop codons, as well as the 3′ UTR occupancy, were consistently higher in the *tma64Δtma20Δ* strain compared to the *tma64Δ* strain, and the same was true when comparing *tma20Δ* and *tma22Δ* single mutants. In some cases, an increase in reinitiation rate was observed in *tma20Δ* but not in *tma22Δ* (Gaikwad et al, 2021), although in some other cases, the effects of these two mutations were equal (Jendruchova et al, 2024). We also draw attention to the fact that *TMA20*, but not *TMA22*, has a statistically significant negative genetic interaction with *TMA64* in genome-wide screenings (Costanzo et al, 2016) and that the *tma20Δ* strain, but not *tma22Δ*, exhibits halfmers in the polysome profile (Fleischer et al, 2006). The latter feature could be explained by a larger defect in either ribosome recycling or biogenesis (Fleischer et al, 2006; Young et al, 2018; Young et al, 2021) in the *tma20Δ* strain. It should also be noted that such analyses are complicated by the mutual dependence of the two proteins on each other for protein stability, as has been previously shown for human and fly cells (see (Ahmed et al, 2018) and references therein). This interdependence is further discussed below.

Although these differential effects of Tma20p and Tma22p depletion were not strongly pronounced, this suggests that Tma22p is not an indispensable component of the heterodimer. This hypothesis aligns well with our results obtained with the Tma22p variant lacking the SUI1 domain but retaining the ZnBD domain (L95*). When produced in *tma64Δtma22Δ* cells, this protein reversed suppression of aberrant reinitiation, albeit to a lesser extent than the *wt* Tma22p protein. This could be explained by a putative ability of Tma20p to promiscuously operate with other SUI1-domain-containing proteins, e.g., Sui1p/eIF1. In this regard, the absence of this rescue effect in the parental *tma64Δtma22Δ* strain could be explained by the previously mentioned degradation of Tma20p when its interacting partner Tma22p is absent. This degradation is likely due to a destabilizing solvent-exposed hydrophobic patch that is normally shielded by the ZnBD domain (Ahmed et al, 2018; Lomakin et al, 2019).

Our hypothesis is further supported by the data obtained with a chimeric protein where the N-terminal ZnBD domain of Tma22p was fused to the SUI1 domain of Sui1p. Production of this protein in the *tma22Δ* strain partially restored reinitiation suppression, indicating the principal ability of the eIF1 SUI1 domain to replace its counterpart in the recycling factor.

Interestingly, in early studies performed in a purified mammalian system, Pestova’s group reported that eIF1 (together with eIF1A and eIF3) can induce tRNA/mRNA release from 40S subunits under specific conditions (Pisarev et al, 2007; Pisarev et al, 2010; Skabkin et al, 2013). In addition, eIF1 has been documented to discriminate against non-initiator tRNAs in the P site of the 40S subunit, effectively removing elongator tRNAs from ribosomal complexes – an activity similar to that needed for post-termination 40S ribosome recycling (Lomakin et al, 2006). Moreover, on an exotic model mRNA (ΔII CSFV IRES) that formed pre-initiation complexes *via* eIF2D- or MCTS1/DENR-dependent delivery of Met-tRNA_i_ in an artificial reconstituted system, eIF1 successfully replaced DENR, acting as MCTS1’s partner in positioning the tRNA on the ribosome (Skabkin et al, 2010). This supports the idea of common principles governed by the SUI1 structural scaffold at distinct steps of the translation cycle (Lomakin et al, 2020; Lomakin et al, 2006).

Our work also contributes to advancing the understanding of the controversial question of how defects in ribosome recycling factors actually impact reinitiation efficiency. As described in detail above, there is a discrepancy in this regard between findings made in animal (Bohlen et al, 2020; Castelo-Szekely et al, 2019; Chen et al, 2022; Haas et al, 2016; Meurs et al, 2025; Schleich et al, 2017; Schleich et al, 2014; Vasudevan et al, 2020) and yeast (Gaikwad et al, 2021; Jendruchova et al, 2024; Makeeva et al, 2019; Young et al, 2018; Young et al, 2021) systems. Although both of them show similar signs of impaired 40S recycling (elevated ribosome occupancy near stop codons and at 3′ UTRs) (Bohlen et al, 2020; Gaikwad et al, 2021; Young et al, 2018; Young et al, 2021), analysis of individual native and reporter mRNAs led to distinct conclusions.

This has been usually attributed to peculiarities of 40S/tRNA/mRNA complex disassembly and propensity of 40S retention on mRNA, and their permissiveness for reinitiation in different systems (discussed in (Bohlen et al, 2020; Gaikwad et al, 2021; Jendruchova et al, 2024; Young et al, 2021)). However, we would also like to draw attention to the differences in the 5′ UTRs and uORFs of the analyzed transcripts. In particular, most of the *Drosophila* and mammalian mRNAs shown to date to depend on MCTS1, DENR, or eIF2D – both natural and reporter transcripts – contained very short uORF(s) within relatively long 5′ UTRs (Bohlen et al, 2020; Castelo-Szekely et al, 2019; Chen et al, 2022; Haas et al, 2016; Meurs et al, 2025; Schleich et al, 2017; Schleich et al, 2014; Vasudevan et al, 2020). Such 5′ UTRs may possess additional features, such as cryptic uORFs arising from non-AUG start codons, complex secondary structure, or protein binding sites, which may significantly complicate interpretation. Strikingly, the dependence on these factors showed a sharp correlation with tiny uORF length, with the most prominent effects observed in human cells featuring only the “zero-length” AUG-stop uORFs (start-stop uORF, uSt-stORF). Such uSt-stORFs are unique, as their translation may involve initiation immediately followed by termination, with the initiator Met-tRNA_i_ positioned in the A site and no peptide in the peptide tunnel (Rendleman et al, 2026). It should also be noted that after 60S subunit and deacylated tRNA dissociation, an mRNA-bound 40S subunit remains with the AUG codon positioned in the P site. This situation is also observed with mRNAs employing non-canonical initiation pathways, which have previously been shown to form 48S pre-initiation complexes with eIF2D, MCTS1/DENR, or eIF5B (reviewed in (Akulich et al, 2016)). This complex formation potentially leads to repetitive cycles of futile initiation-termination-splitting-rejoining (ITSR). Furthermore, in the absence of MCTS1/DENR, eIF1 can easily enter such a complex, but it likely may not be able to remove the tRNA, unlike in a canonical post-termination complex, because its ability to discriminate between initiator and elongator tRNAs may not function properly in this context (Lomakin et al, 2006).

In contrast, yeast mRNAs usually have relatively short and simple 5′ UTRs. We previously used luciferase reporters with a CA-rich leader, modeling reinitiation in both the 5′ and 3′ UTRs, to dissect the effects of Tma protein depletion in yeast cell-free systems (Young et al, 2018). Our new *in vivo* data were obtained with similar reporters (uORF_rein and mORF_rein). However, our results align well with the recent study using *LacZ* reporters with a relatively long 5′ UTR derived from the native *GCN4* leader (Jendruchova et al, 2024). We also note that the penultimate codons in our uORF_rein and mORF_rein constructs are CAA^Gln^ and UUA^Leu^, respectively. CAA^Gln^ was considered to render both uORF and mORF “DENR-independent” in mammals (Bohlen et al, 2020) and yeast (Young et al, 2021), while UUA^Leu^ is considered “DENR-neutral”. In a *GCN4*-derived reporter, CAA^Gln^ was shown to produce a 1.4-fold and a 2.4-fold increase in reinitiation rate in *tma20Δtma22Δ* and *tma20Δtma22Δtma64Δ* strains, respectively, compared to the *wt* strain (Jendruchova et al, 2024). This corresponds to the 6.8-fold and 11.3-fold increases we observed in our uORF_rein construct, which had the same penultimate codon. The more pronounced effect in our case is likely due to the fact that in our reporter, the uORF directly abutted the CDS, whereas in the colleagues’ study, it was spaced further away. As we previously demonstrated *in vitro*, spacing the uORF reduces the effects of *TMA* gene mutations on reinitiation suppression (Young et al, 2018).

Nevertheless, we can see that in the case of truly complex 5′ UTRs (not only long but also containing complicated uORF systems), such as the *wt GCN4* leader (Gaikwad et al, 2021; Jendruchova et al, 2024; Makeeva et al, 2019), as well as in the case of reinitiation-permissive uORFs or uSt-stORFs (Jendruchova et al, 2024), discordant effects from Tma protein depletion can be observed. This may mean that on long and complex 5′ UTRs, a composite cross-talk between multiple translation rounds of AUG- and non-AUG-initiated uORFs, scanning, and reinitiation can occur, making it challenging to dissect the specific contribution of each component to the overall molecular mechanism.

MCTS1, DENR, and eIF2D have been implicated in numerous human pathological states, including cancer, immune deficiencies, autism, schizophrenia, and other neurological disorders (Grove et al, 2024; Shyrokova et al, 2021; Zamyatnina, 2025). Specifically, mutations in the human *DENR* gene resulting in amino acid substitutions C37Y and P121L are associated with autism spectrum disorder and Asperger syndrome (Haas et al, 2016; Neale et al, 2012). C37 is located in the ZnBD and forms the zinc-binding pocket (Ahmed et al, 2018; Lomakin et al, 2019). P121 is the amino acid residue preceding the first conserved Arg of the basic loop studied here and likely contributes to local rigidity, thereby enhancing the loop’s activity in the dissociation of the P-site tRNA (Lomakin et al, 2020; Lomakin et al, 2017; Weisser et al, 2017). Orthologous substitutions (C11Y and A105L) in yeast Tma22p were recently analyzed for their effects on translation reinitiation using TCP-seq (Young et al, 2021). While C11Y did not rescue the 40S recycling defect, which aligns well with our own observations with strains lacking the ZnBD (L2-E90del), Tma22p-A105L proved as active as the *wt* protein. This raised the question of the importance of the conserved β-hairpin loop 1 for DENR/Tma22p function. However, in yeast eIF1 (Sui1p), the effects of perturbations in this structural element are well documented (Martin-Marcos et al, 2013). Substitutions of Sui1p residues R33 and K37 (corresponding to R106 and K110 of Tma22p) with Glu are lethal, while other substitutions in this region (e.g., R33A, N34E, R36G, R36A, R36E, and K37A) confer a slow growth phenotype, increased initiation on UUG codons, or both. *In vitro*, R33A and K37E weakened Sui1p binding to the 40S subunit. The basic loop of human DENR is positioned on the 40S P site (Lomakin et al, 2017; Weisser et al, 2017) similarly to that of eIF1 (Hussain et al, 2014; Llacer et al, 2015; Lomakin et al, 2003; Lomakin & Steitz, 2013; Rabl et al, 2011; Weisser et al, 2013). This loop protrudes toward the mRNA binding channel in the P site, with residues K125 and K126 (corresponding to R109 and K110 in yeast Tma22p, respectively) contacting helix h44 of the 18S rRNA (Lomakin et al, 2017). In our study, we show that two Tma22p variants, either with a single substitution K110E or with a larger modification KRK108AEA of loop 1, are completely unable to rescue the *tma64Δtma22Δ* phenotype. Thus, the positively charged β-hairpin loop 1 of the Tma22p SUI1 domain is a critical determinant of its function. These data may be important for the diagnosis of hereditary neurological diseases.

In conclusion, our results highlight a striking functional parallelism between initiation and recycling machineries, where a common SUI1 structural scaffold has been evolutionarily co-opted to regulate distinct steps of the translation cycle.

## Materials and Methods

### Reporter plasmids

All oligos used for cloning and verification are listed in Supplementary Table S1. To prepare the uORF_rein and Control*_*1 plasmids, the *Fluc* CDS was first amplified from the pGL3R plasmid (Promega) by PCR using primers BamHI_Fluc*_*fw and PstI_Fluc*_*rev. After digestion with BamHI and PstI, it was inserted into the corresponding sites of the pCM189 vector (a gift from Alexander Alexandrov) under the control of the pBAD/tetO promoter, yielding pCM189-Fluc1. The *Nluc* gene was amplified from the pNL1.1 plasmid (Promega) using either primer pair BamHI_uORF*_*Nluc_fw and PstI*_*Nluc_rev, or BamHI*_*noORF_Nluc*_*fw and PstI_Nluc*_*rev. After digestion with BamHI and PstI, it was inserted into the same sites of pCM189 to yield pCM189-uORF-Nluc and pCM189-no-uORF-Nluc, respectively. Next, the *Nluc* expression cassettes were amplified from these two plasmids using primers HindIII_ADH1term*_*fw and HindIII_CYC1term*_*rev. After digestion with HindIII, they were inserted into pCM189-Fluc1 to yield the final uORF_rein and Control*_*1 plasmids, respectively.

To create the mORF_rein plasmid, the *Fluc* CDS was amplified from pGL3R using primers BamHI_Fluc_fw and PstI_AgeI_Fluc_rev. After digestion with BamHI and PstI, it was inserted into pCM189, yielding pCM189-Fluc2. Subsequently, the *Nluc* gene was amplified from the pNL1.1 plasmid (Promega) using primers AgeI_Nluc_rein_fw and PstI_Nluc_rev. Following digestion with AgeI and PstI, the *Nluc* fragment was inserted between the corresponding sites of pCM189-Fluc2 to yield mORF_rein. The Control_2 plasmid was constructed similarly. First, Fluc was amplified from pGL3R using BamHI_Fluc_fw and PstI_AgeI_P2A_Fluc_rev primers. After digestion with BamHI and PstI, it was inserted into pCM189, yielding pCM189-Fluc-P2A. Subsequently, Nluc was amplified from pNL1.1 using AgeI_Nluc_fw and PstI_Nluc_rev primers. Following digestion with AgeI and PstI, the Nluc fragment was inserted between the same sites of pCM189-Fluc-P2A to yield the final Control_2 plasmid.

### Yeast strain construction and TMA22 mutagenesis

Yeast strains used in this study are listed in Supplementary Table S2. All strains were either in the BY4741 or W303 background, as indicated. The primers used for plasmid and strain construction, verification, and mutagenesis are listed in Supplementary Table S1.

The *tma20Δ*, *tma22Δ*, *tma64Δ*, *tma64Δtma20Δ*, and *tma64Δtma22Δ* strains were either from Dharmakon or described previously (Young et al, 2018). To generate the *tma20Δtma22Δ* strain, a one-step gene disruption approach was used. A disruption cassette containing the *HIS3* selectable marker was amplified by PCR using the pRS313 vector (ATCC) as a template. Primers TMA20-HIS3-F and TMA20-HIS3-R were designed to flank the *TMA20* locus with homology arms of ∼50 bp on each side. The resulting PCR product was used to transform the parental *tma22Δ* strain. Transformants were selected on synthetic complete medium lacking histidine (SC-His). Correct integration of the *HIS3* cassette at the *TMA20* locus was confirmed by colony PCR. The resulting strain was designated LSBS001 (*tma22Δtma20Δ*). The same *HIS3*-containing PCR product was similarly used to transform the *tma64Δtma22Δ* strain, resulting in LSBS002 (*tma64Δtma22Δtma20Δ*).

The *TMA22* CDS from *S. cerevisiae* BY4741 cDNA was amplified by PCR with primers Tma22_for and Tma22_rev and cloned into pET11c (Novagen) using NdeI and BamHI sites. Concurrently, the *SUI1* CDS, amplified with F_eIF1 and R_eIF1_HindII, was cloned into pET22b(+) (Novagen) using NdeI and HindIII sites. The resulting plasmids were named pET11c-TMA22 and pET22b(+)-SUI1, respectively. To prepare the pET11c-TMA22 variant encoding the Tma22p-K110E protein, site-directed mutagenesis was performed by inverse PCR of the full plasmid using primers F_K110E_TMA22 and R_K110E_TMA22.

Next, the *TMA22_WT_His3* and *TMA22_K110E_His3* cassettes were obtained by overlap extension PCR. First, the native or mutated *TMA22* gene was amplified from the corresponding plasmids (pET11c-TMA22 and pET11c-TMA22-K110E, respectively) using primers TMA22_fw and TMA22*_*rv. The *HIS3* selectable marker was amplified from pFA6a-3HA-His3MX6 (Addgene plasmid #41600 (Longtine et al, 1998)) using primers ADH1term_fw and TEFterm_rv. Subsequently, overlap extension PCR was performed using the amplified *TMA22* gene fragments and the *HIS3* marker fragment, with primers TMA22_fw and TEFterm_rv to join them. Then, the assembled *TMA22_K110E_His3* cassette was amplified using primers TMA22cassDir and TMA22cassRev to generate the final integrative fragment. The resulting PCR product was purified and used to transform the yeast strain YDY12 (*tma64Δtma22Δ*). Transformants were selected on SC-His medium and verified by colony PCR. The resulting strain carrying the *TMA22_K110E* allele was designated LSBS004 (*tma64Δtma22Δtma22-K110E*).

To create the strain expressing the KRK108-110AEA derivative of the Tma22p protein, two DNA fragments of the *TMA22_WT_His3* gene cassette were amplified using primer pairs TMA22cassDir and TMA22-108AEA_Rev (for the N-terminal fragment) and TMA22cassRev and TMA22-108AEA_Dir (for the C-terminal one). The resulting fragments were then used as templates for overlap extension PCR with the primer pair TMA22cassDir and TMA22cassRev to assemble the full-length product. The assembled cassette, designated *TMA22_108AEA_His3*, was used to transform the YDY12 strain, yielding the LSBS003 strain (*tma64Δtma22Δtma22-K108A_R109E_K110A*).

To create the strain producing the L95* Tma22p variant with a deletion of the SUI1 domain, we introduced a stop codon at the corresponding position in the *TMA22* gene. Two DNA fragments were amplified from the *TMA22_WT_His3* gene cassette using primer pairs TMA22cassDir and TMA22-95X_Rev (to amplify the N-terminal fragment) and TMA22cassRev and TMA22-95X_Dir (to amplify downstream sequence), creating a truncated *TMA22* allele. The resulting fragments were then used as templates for overlap extension PCR with the primer pair TMA22cassDir and TMA22cassRev to assemble the full-length product. The assembled cassette, designated *TMA22_95*_His3,* was purified and used to transform the yeast strain YDY12, followed by selection on SC-His medium and verification by colony PCR. The resulting strain carrying the *TMA22_95*_His3* allele was designated LSBS005 (*tma64Δtma22Δtma22-95\**). The strain producing the L2_E90del Tma22p derivative lacking the N-terminal domain was created similarly. For this, two DNA fragments were amplified from the *TMA22_WT_His3* gene cassette using primer pairs TMA22cassDir and TMA22-d2-90_Rev (for the N-terminal fragment) and TMA22cassRev and TMA22-d2-90_Dir (for the remaining segment). The resulting fragments were then used as templates for overlap extension PCR with the primer pair TMA22cassDir and TMA22cassRev to assemble the full-length product. The assembled cassette, designated as *TMA22_L2-E90del_His3*, was used to transform the YDY12 strain, yielding LSBS006 (*tma64Δtma22Δtma22-L2-E90del*).

To create a construct encoding the Tma22p-Sui1p chimera (consisting of amino acid residues 1–77 of Tma22p and residues 27–108 of Sui1p), plasmids pET11c-TMA22 and pET22b(+)-SUI1 were used as templates. Two PCR reactions were performed: one using primers TMA22_rev (1-75) and R_chimera (to amplify the *TMA22* fragment), and the other using primers F_chimera and R_eIF1_HindIII (to amplify the *SUI1* fragment). F_chimera and R_chimera are complementary to each other, facilitating the fusion of the two PCR products in a subsequent PCR step using primers TMA22_rev (1-75) and R_eIF1_HindIII. The resulting *tma22_sui1* PCR product was digested with NdeI and HindIII and cloned into the corresponding sites of pET22b(+), yielding pET22-Chimera. Next, the chimeric gene was amplified from pET22-Chimera using primers ChimYIplac_F2.0 and ChimYIplac_R2.0, digested with HindIII and NdeI, and cloned into the corresponding sites of YIplac128 (gifted by Svyatoslav Sokolov), resulting in pChim-YIplac128. Subsequently, the promoter region of the *TMA22* gene was amplified from genomic DNA of the BY4741 *wt* strain with primers Prom22Hind_F and Prom22Hind_R, digested with HindIII, and inserted into the HindIII site of pChim-YIplac128 to yield the integrative pProm_Chim_YIplac128 plasmid.

To generate the W303 strain expressing the chimera, the parental W303 *wt* strain (gifted by Svyatoslav Sokolov) was transformed with pProm_Chim_YIplac128 linearized by digestion with ClaI. Transformants with the plasmid integrated into the targeted *leu2-3* locus were selected on synthetic complete medium lacking leucine (SC-Leu). The resulting strain was designated KAZ001 (*Prom_Chim_LEU2*).

To generate a *TMA22* deletion in the W303 *wt* and KAZ001 strains, a disruption cassette containing the *kanMX* marker was amplified by PCR using genomic DNA from the 6812 (*tma22Δ*) strain as a template, with primers TMA22_A and TMA22D2, producing homology arms of ∼200 bp on each side. The parental W303 *wt* and KAZ001 strains were transformed with this product. Transformants were selected on YPD plates containing 250 μg/mL G418, yielding deletion strains KAZ002 (*tma22Δ*) and KAZ003 (*Prom_Chim_LEU2 tma22Δ*). Correct integration was confirmed by colony PCR using primers TMA22_A and Kan_B, with no product detected when using primers TMA22_A and TMA22_B.

### Yeast growth, cell lysis, and luciferase assays

Yeast transformations were performed using the LiOAc method. Transformants were selected on appropriate synthetic complete media (0.17% yeast nitrogen base without amino acids, 0.5% ammonium sulfate, 2% glucose, supplemented with required amino acids) and incubated at 30 °C for 2-3 days. Single colonies were inoculated into 2 mL of selective liquid medium and grown overnight at 30 °C with shaking at 220 rpm. The next day, cultures were diluted to an OD_600_ of 0.3 in 5 mL of standard YPD medium (1% yeast extract, 2% peptone, 2% glucose) and grown to OD_600_ 0.6–1.0. Cells were harvested by centrifugation, washed with deionized water, and lysed in 100 μL of 1× Passive Lysis Buffer (Promega) by gentle rocking at room temperature for 15 min. Lysates were cleared by centrifugation at 14,000 rpm, and the supernatants were used for luciferase assays.

Luciferase activities were measured using 2.5 μL of lysate and the Dual Luciferase Reporter Assay System (Promega). Reactions were performed sequentially with 10 μL of LARII (for Fluc) and then 10 μL of SG reagent (for Nluc) using a TD-20/20 Compact Luminometer (Turner Designs Instruments) with a 10 s signal integration time.

### Statistical analyses

Translation reinitiation efficiency was quantified as the ratio of Nluc to Fluc activity measured in each independent yeast clone. All Nluc/Fluc values were log₂-transformed prior to analysis. For each mutant, the reinitiation rate was assessed as the difference (Δ) between the mean log₂(Nluc/Fluc) of the reporter construct carrying the uORF or mORF and the mean log₂(Nluc/Fluc) of the matched “Control” construct. The standard error of this difference was calculated by error propagation under the assumption of independent samples: SE(Δ) = √(S²_reporter_ / n_reporter_ + S²_control_ / n_control_), where S² denotes the sample variance and n is the number of biological replicates for each construct within a given genotype. 95% confidence intervals were calculated as Δ ± 1.96 × SE(Δ). To express mutant reinitiation relative to *wt*, the *wt* log₂ score was subtracted from each mutant log₂ score.

Pairwise differences between all mutants were assessed using a two-tailed z-test applied to the difference of log₂ effect estimates, with the combined standard error SE(diff) = √(SE²_G1_ + SE²_G2_) as the denominator (where G1 and G2 denote different mutant genotypes). The resulting *p*-values were adjusted for multiple comparisons using the Holm-Bonferroni step-down procedure. Mutant pairs with an adjusted *p*-value below 0.001 were considered statistically significant. All analyses were performed in Python (v.3.x) using the pandas, NumPy, and SciPy libraries; multiple-testing correction was applied with the multipletests function from the statsmodels package.

## Supporting information

Supplementary Data - Plasmid Maps

## Abbreviations

eIF: eukaryotic initiation factor;
CDS: coding sequence; Fluc, firefly luciferase;
KO: knockout; Nluc, Nano-luciferase;
ORF: open reading frame;
UTR: untranslated region;
*Wt*: wild type

## Author Contributions

S.E.D. and K.A.Z. conceived the study and designed the experiments. I.A.V., E.A.S., and K.A.Z. assembled genetic constructs and DNA vectors. V.N.U., V.V.K., and K.A.Z. constructed and provided yeast strains. K.A.Z. conducted experiments. E.S.G., L.M.K., and K.A.Z. performed statistical analysis. K.A. Z., S.E.D., and P.A.K. interpreted the data. K.A.Z. and S.E.D. drafted the manuscript and prepared the figures. S.E.D., P.A.K., and I.A.V. contributed to editing the manuscript. S.E.D. supervised the study.

## Acknowledgments

We are grateful to Ivan Lomakin for seminal discussions and invaluable suggestions at the initial stage of the study. We thank Ivan Chicherin, Svyatoslav Sokolov, Alexander Alexandrov, and Anastasia Sukhinina for their experimental advice and suggestions, the provision of the W303 *wt* strain, YIplac128, and pCM189 plasmids, and for discussing the manuscript.

