## Supplementary Table and Figures for "The conserved β-hairpin of the SUI1 domain is a dual-function structural module governing translation initiation and ribosome recycling in yeast"

---

### Supplementary Table S1. DNA oligonucleotides used for cloning and strain construction

| Oligonucleotide | Nucleotide sequence (5' to 3') |
|---|---|
| Tma22_for | GGCCCATATGTTAAGAGAAGTCATCTATTG |
| Tma22_rev | GCCGGGATCCTTACTTGGCAGCTCCTTCTG |
| F_K110E_TMA22 | GCTAGGACCAAGAGAGAATTTATCGTCGCTATC |
| R_K110E_TMA22 | GATAGCGACGATAAATTCTCTCTTGGTCCTAGC |
| F_eIF1 | GGCCCATATGTCCATTGAGAATCTGAAATC |
| R_eIF1_HindIII | GCCGAAGCTTTAAAACCCATGAATTTTAATG |
| TMA22rev (1-75) | GCCGGGATCCTTAGTCCTTTTCCAACTTTTC |
| F_chimera | GGAAAAGGACTTGCTGCATATTCGTATCCAAC |
| R_chimera | GTTGGATACGAATATGCAGCAAGTCCTTTTCC |
| TMA20-HIS3-F | ATATAAAAGACAAAGACGGTAAACTAAAACAGCAGAGAGGAACGTTTTAAGAGCTTGGTGAGC |
| TMA20-HIS3-R | TGTAGCAGATGGATAGTAATATAGTGTTGACGGCTCCGTTTGTTATCGAGTTCAAGAGAAAAAAAAAGAA |
| TMA22cassDir | CCCAAGGAAACAGTTCAAGAG |
| TMA22-108AEA_Rev | AAGCTTCAGCGGTCCTAGCTTCTCTTTTAATGA |
| TMA22cassRev | TAAAAAGTCCTTTTCTCCCAGAAC |
| TMA22-108AEA_Dir | GCTAGGACCGCTGAAGCTTTTATCGTCGCTATCTCTGGG |
| TMA22-95X_Dir | GCTAAGAAGTAAGCAATCCATAAATATGTAATAGCA |
| TMA22-95X_Rev | TATTTATGGATTGCTTACTTCTTAGCTAATTCTCTTTGTTCC |
| TMA22-d2-90_Rev | GCTTCTTAGCTAACATTTTATGCAATATGCTTTTCT |
| TMA22-d2-90_Dir | GCATAAAATGTTAGCTAAGAAGCTGTCATCGAA |
| ChimYIplac_F2.0 | ACCAAAAGCTTAACAAAATGTTAAGAGAAGTCATCTATTGTGG |
| ChimYIplac_R2.0 | CCACACATATGTTAAAACCCATGAATTTTAATGTTC |
| Prom22Hind_F | CCACAAAGCTTACTGACGTGAACTTGCACCATG |
| Prom22Hind_R | CCACAAAGCTTTTTATGCAATATGCTTTTCTTTAGTTT |
| TMA22_A | CTGACGTGAACTTGCACCATG |
| TMA22D2 | GAAGGTTTCAGAGAAGGAACG |
| Kan_B | GTATATGAAAGAAGAACCTCAGTG |
| TMA22_B | TTTTCAGCAAGTCCTTTTCCAAC |
| BamHI_Fluc_fw | GCAACGGATCCAACAACAACAACAACAACAAAATGGAAGACGCCAAAAACATAAAG |
| PstI_AgeI_Fluc_rev | GCAACCTGCAGGCATACACCGGTCACGGCGATCTTTCCGC |
| AgeI_Nluc_rein_fw | GCAACACCGGTGAATTATGACTGTCTTCACACTCGAAGATTTCGTT |
| PstI_Nluc_rev | GCAACCTGCAGTTACGCCAGAATGCGTTCG |
| PstI_AgeI_P2A_Fluc_rev | GCAACCTGCAGGCATACACCGGT**TGGACCTGGATTCAATTCAACATCACCAGCCAATTTCAACAAAGAAAAATTAGTAGCACCACCAGAACCAGT**CACGGCGATCTTTCCGC |
| AgeI_Nluc_fw | GCAACACCGGTATGACTGTCTTCACACTCGAAGATTTCGTT |
| PstI_Fluc_rev | GCCCTGCAGTTACACGGCGATCTTTCCG |
| BamHI_uORF_Nluc_fw | GCCGGATCCAACAACAACAACAACAAAATGACACAAACACAATGACTGTCTTCACACTCGAAGATTTCGTT |
| BamHI_noORF_Nluc_fw | GCCGGATCCAACAACAACAACAACAAAAGTACACAAACACAATGACTGTCTTCACACTCGAAGATTTCGTT |
| HindIII_ADH1term_fw | GCAACAAGCTTATAAGCGAATTTCTTATGATTTATGATTTTTATTATT |

| | |
|---|---|
| HindIII_CYC1term_rev | GCAACAAGCTTGGCCGCAAATTAAA |
| TMA22_fw | CCCAAGGAAACAGTTCAAGAGCTAAACTAAAGAAAAGCATATTGCATAAAATGTTAAGAGAAGTCATCTATTGTGGA |
| TMA22_rv | GAAATTCGCTATGGATTGCTTACTTGGCAGCTCCTTCTG |
| ADH1term_fw | AGCAATCCATAGCGAATTTCTTATGATTTATGATTTTTATTATTAAAT |
| TEFterm_rv | TAAAAAGTCCTTTTCTCCCAGAACGGTGCTATTACATATTTATGGATTGCCAGTATAGCGACCAGCATTCA |

Restriction sites are underlined. The sequence corresponding to the 2A peptide from Equine rhinitis B virus is marked in bold.

**Supplementary Table S2. Yeast strains used in this study**

| Name | Genotype | Source |
|---|---|---|
| BY4741 | *MATa his3Δ1 leu2Δ0 met15Δ0 ura3Δ0* | Dharmacon |
| 328 | *MATa his3Δ1 leu2Δ0 met15Δ0 ura3Δ0 tma20Δ::KanMX4* | Dharmacon |
| *6812* | *MATa his3Δ1 leu2Δ0 met15Δ0 ura3Δ0 tma22Δ::KanMX4* | Dharmacon |
| FZY793 | *MATa his3Δ1 leu2Δ0 met15Δ0 ura3Δ0 tma64Δ::HygMX4* | Young et al., 2018 |
| YDY10 | *MATa his3Δ1 leu2Δ0 met15Δ0 ura3Δ0 tma64Δ::HygMX4 tma20Δ::KanMX4* | Young et al., 2018 |
| YDY12 | *MATa his3Δ1 leu2Δ0 met15Δ0 ura3Δ0 tma64Δ::HygMX4 tma22Δ::KanMX4* | Young et al., 2018 |
| LSBS001 | *MATa his3Δ1 leu2Δ0 met15Δ0 ura3Δ0 tma22Δ::KanMX4 tma20::His3* | This study |
| LSBS002 | *MATa his3Δ1 leu2Δ0 met15Δ0 ura3Δ0 tma64Δ::HygMX4 tma22Δ::KanMX4 tma20::His3* | This study |
| LSBS003 | *MATa his3Δ1 leu2Δ0 met15Δ0 ura3Δ0 tma64Δ::HygMX4 tma22Δ::KanMX4 tma22-K108A_R109E_K110A* | This study |
| LSBS004 | *MATa his3Δ1 leu2Δ0 met15Δ0 ura3Δ0 tma64Δ::HygMX4 tma22Δ::KanMX4 tma22-K110E* | This study |
| LSBS005 | *MATa his3Δ1 leu2Δ0 met15Δ0 ura3Δ0 tma64Δ::HygMX4 tma22Δ::KanMX4 tma22-95** | This study |
| LSBS006 | *MATa his3Δ1 leu2Δ0 met15Δ0 ura3Δ0 tma64Δ::HygMX4 tma22Δ::KanMX4 tma22-L2-E90del* | This study |
| W303 | *MATa/MATα {leu2-3,112 trp1-1 can1-100 ura3-1 ade2-1 his3-11,15} [phi+]* | A gift from Svyatoslav Sokolov |
| KAZ001 | *MATa/MATα {leu2-3,112 trp1-1 can1-100 ura3-1 ade2-1 his3-11,15} [phi+], tma22-sui1* | This study |
| KAZ002 | *MATa/MATα {leu2-3,112 trp1-1 can1-100 ura3-1 ade2-1 his3-11,15} [phi+] tma22Δ :: KanMX4,* | This study |
| KAZ003 | *MATa/MATα {leu2-3,112 trp1-1 can1-100 ura3-1 ade2-1 his3-11,15} [phi+] tma22Δ :: KanMX4, tma22-sui1* | This study |

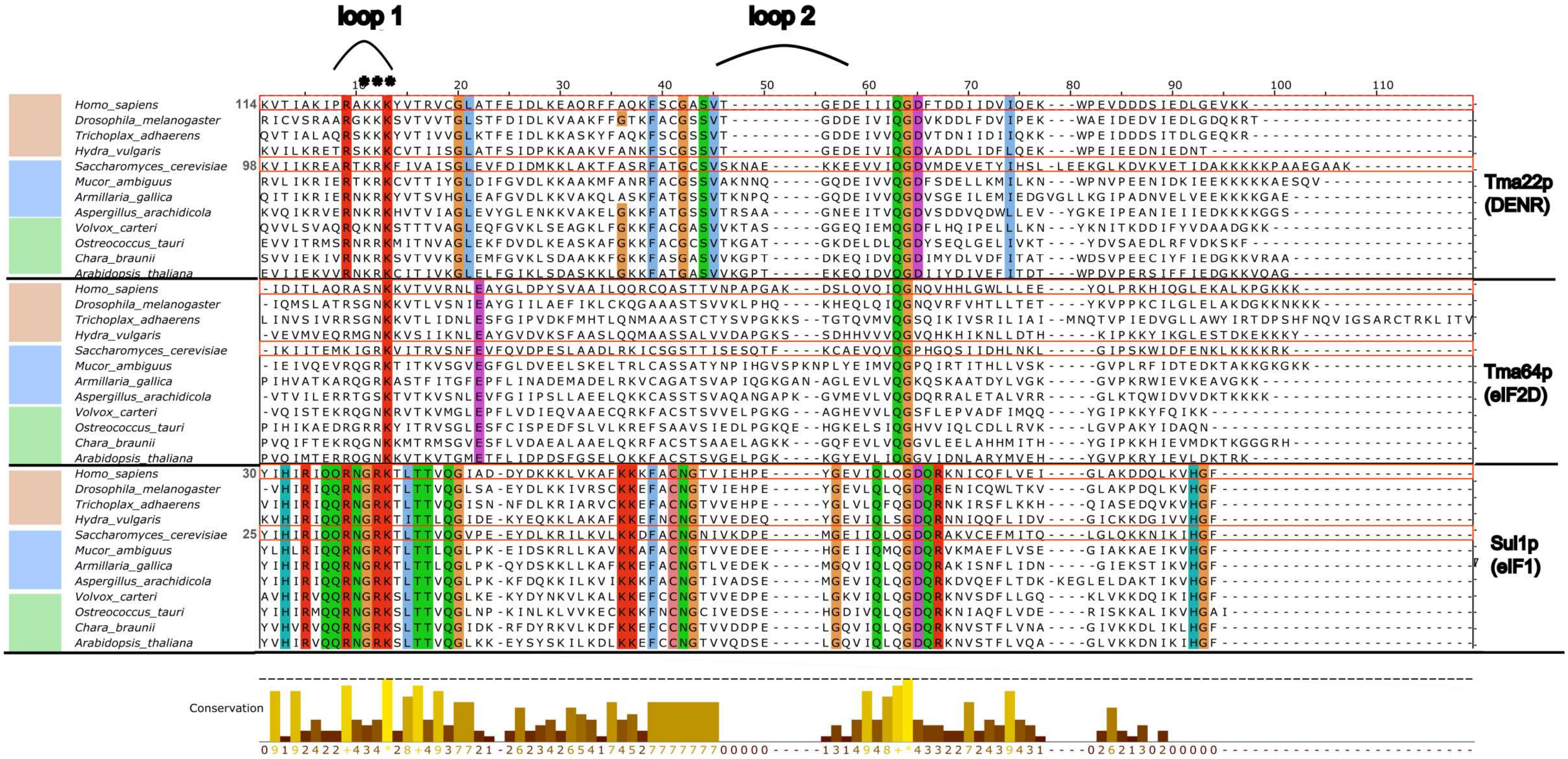


**Supplementary Figure S1.** Alignment of amino acid sequences of SUI1 domains from Tma22p/DENR, Tma64p/eIF2D, and Sui1p/eIF1 proteins from a representative set of eukaryotic organisms. Animal, fungi, and plant species are color-coded. For Tma22p/DENR and Sui1p/eIF1 from human and budding yeast, starting residue numbers are shown. Conserved residues are highlighted according to amino acid properties, and the positions of β-hairpin loops 1 and 2 are marked above the alignment. Alignment was generated using ESPript 3.
